# Parasite-mediated inbreeding depression in wild red deer

**DOI:** 10.1101/2025.02.06.636807

**Authors:** Adam Z. Hasik, Anna M. Hewett, Katie Maris, Sean J. Morris, Ali Morris, Gregory F. Albery, Josephine M. Pemberton

**Author notes:** **Author for correspondence**: Adam Z. Hasik, Institute of Ecology and Evolution, University of Edinburgh, Edinburgh, UK.

## Abstract

Inbreeding depression is the reduction in fitness of inbred individuals relative to their more outbred counterparts. Parasitism also reduces fitness and is a route by which inbreeding depression may operate, yet the complete pathway from inbreeding to parasitism to fitness has almost never been documented in the wild. Using high-quality, individual-level data on fitness in juveniles and adult females, longitudinal infection data for three gastrointestinal helminth parasites, and genomic inbreeding coefficients we test for parasite-mediated inbreeding depression in a wild ungulate population (red deer, *Cervus elaphus*). We found evidence for parasite-mediated inbreeding depression via strongyle nematodes in juvenile survival, independent of direct adverse effects of inbreeding on survival and indirect effects of inbreeding on survival via birth weight. Inbreeding also reduced fitness in reproductive adults by reducing overwinter survival. Our study reveals three independent pathways by which inbreeding depresses fitness and highlights the rarely-studied route of parasitism.

## Introduction

Inbreeding depression, the reduction in fitness in the offspring of related parents, is a universal phenomenon of diploids. It is ascribed to two mechanisms: the expression of (partially) deleterious alleles, generally thought to be the more important process, and loss of heterozygosity at loci with heterozygote advantage (overdominance) (1). Although we have known about inbreeding depression since Darwin (2), its study in natural populations with biparental inbreeding (i.e., including most animals) has been relatively slow to develop due to the requirement for both estimates of individual fitness and pedigrees or other tools for assessing individual identity-by-descent (IBD). With the advent of genomic approaches for assessing IBD, it is now clear that inbreeding depression in the wild is often more severe than has been documented in the past (3, 4).

Inbreeding depression specifically refers to fitness. Fitness is a hard-to-measure, complex trait which is the outcome of many aspects of an individual’s development and physiology, and these in turn are determined by a combination of environmental and genetic sources of variation. Traits such as body size are often positively related to fitness components such as survival, and are easier to measure than fitness or its components, so many studies of inbreeding depression in the wild focus on such fitness-related traits. However, inbreeding depression (*sensu stricto*) is likely to be expressed via multiple different pathways and at present we have little idea of their nature and relative importance in inbreeding depression.

Parasitism is a strong selective force in natural populations (5), having led to the evolution of multiple physiological mechanisms of resistance or tolerance, including the entire vertebrate immune system (6, 7). A strong theme within host-parasite coevolution research is the idea that parasites select for host genetic diversity because with short generation times they can evolve faster than hosts, and that hosts with higher genetic diversity can resist more parasite species or parasite genotypes. For example, one hypothesis for the maintenance of sexual reproduction among eukaryotes is parasite-mediated selection for increased genetic diversity (8, 9), and parasitism is often cited as the likely cause of the extreme diversity of the mammal major histocompatibility complex (MHC) (10–12). Parasite defense thus seems a likely path by which inbreeding depression could be expressed, perhaps especially through the overdominance mechanism.

The relationship between inbreeding, or correlates of it, and parasitism in natural populations has been researched in a number natural populations, though often with imprecise tools. Studies of Galápagos hawks (13), mountain white-crowned sparrows (14), and California sea lions (15) have found that parasitism increases with estimates of individual inbreeding. However, these examples only considered the relationships between inbreeding and parasitism. Further, these studies also suffer from the fact that they used relatively crude measures of individual inbreeding. For example, the relationship between heterozygosity at a few microsatellites and inbreeding coefficients from a deep pedigree is often weak (16, 17). With time, there has been an expansion in the size and completeness of genetically-supported wild pedigrees, enabling more precise estimation of pedigree inbreeding coefficients (18). This approach has itself been largely supplanted by our ability to genotype thousands of genome-wide markers (3, 4). Genomic measures of individual inbreeding, such as genome-wide homozygosity (19), metrics derived from the genomic relationship matrix (20), and runs of homozygosity (21) now offer even more precise estimates of individual inbreeding than pedigree inbreeding coefficients. This is because whereas pedigree inbreeding coefficients are the mean expectation of IBD, genomic tools generate the realized IBD that results from the random events of Mendelian segregation and recombination. In consequence, inbreeding metrics derived from genome-wide SNP markers generally find stronger evidence of inbreeding depression than all previous methods (3, 4, 19, 22, 23).

A complete understanding of parasite-mediated inbreeding depression in wild populations requires three individual-based components: accurate estimates of the level of inbreeding (estimated from a pedigree or genomic information), parasitism, and, crucially, fitness. Linking these three components is key, as it may uncover the effects of inbreeding on parasitism, along with any subsequent fitness loss. To date, only one study has simultaneously investigated all three components to study parasite-mediated inbreeding depression, finding that hosts that were homozygous at a panel of microsatellites had more parasites and lower overwinter survival, and that parasitism reduced overwinter survival in Soay sheep (24). These results suggest that parasitism is a route by which inbreeding depression is expressed, but more evidence from other wild host-parasite systems is sorely needed to understand the generality of this pattern.

Our goal for this paper was to test for parasite-mediated inbreeding depression in a wild population. We investigated the links between individual-level inbreeding, parasitism by gastrointestinal helminths, and fitness through spatially-explicit analyses and structural equation models using data across two age classes from an exceptionally well-characterized study system of red deer (*Cervus elaphus*). We focused our analyses on juveniles and adult females, as parasite-mediated effects on survival in this system are apparent in both groups (25–27), and there is also evidence for inbreeding depression manifesting through birth weight, juvenile survival, and the likelihood that an adult female has a calf (28). Linking these relationships in a single study allows us to establish what role, if any, parasites play in these relationships.

## Results

### Inbreeding depresses fitness via multiple, independent pathways

We fit spatially-explicit structural equation models to unravel the relationships between inbreeding, parasitism, and fitness using precise genomic measures of inbreeding (proportion of the genome in runs of homozygosity, F_ROH_), overwinter survival, and individual-level infection data (fecal egg counts on the log scale, lEPG) for three gastrointestinal helminths: strongyle nematodes (hereafter “strongyles”, a mix of different species with indistinguishable eggs), liver fluke (*Fasciola hepatica*), and tissue worms (*Elaphostrongylus cervi*) collected three times a year between 2016 and 2023. Our analyses revealed three routes by which inbreeding depression manifests within juveniles in the population. First, we found that F_ROH_ was negatively associated with birth weight (Fig. 1). Given that birth weight was strongly positively associated with survival in the strongyle dataset (Fig. 1a), this indicates that inbreeding has an indirect negative effect on survival via birth weight. Second, F_ROH_ was strongly and negatively associated with overwinter survival in all three datasets (Fig. 1), confirming previous findings of inbreeding depression in this population (22, 28, 29) and providing a direct pathway by which inbreeding depression manifests. The third and final route by which inbreeding depression manifests provided evidence for parasite-mediated inbreeding depression in the strongyle dataset. That is, F_ROH_ was weakly positively associated with strongyle lEPG, and lEPG was strongly and negatively associated with overwinter survival (Fig. 1a). Note that these three routes by which inbreeding depression manifests (indirect negative relationship with survival via birth weight, direct negative relationship with survival, and indirect negative relationship via strongyle lEPG) are independent of one another. lEPG was also weakly and positively associated with F_ROH_ in the *E. cervi* dataset, though there were no relationships between lEPG and survival in the *F. hepatica* or *E. cervi* datasets (Fig. 1b,c).

**Figure 1.**
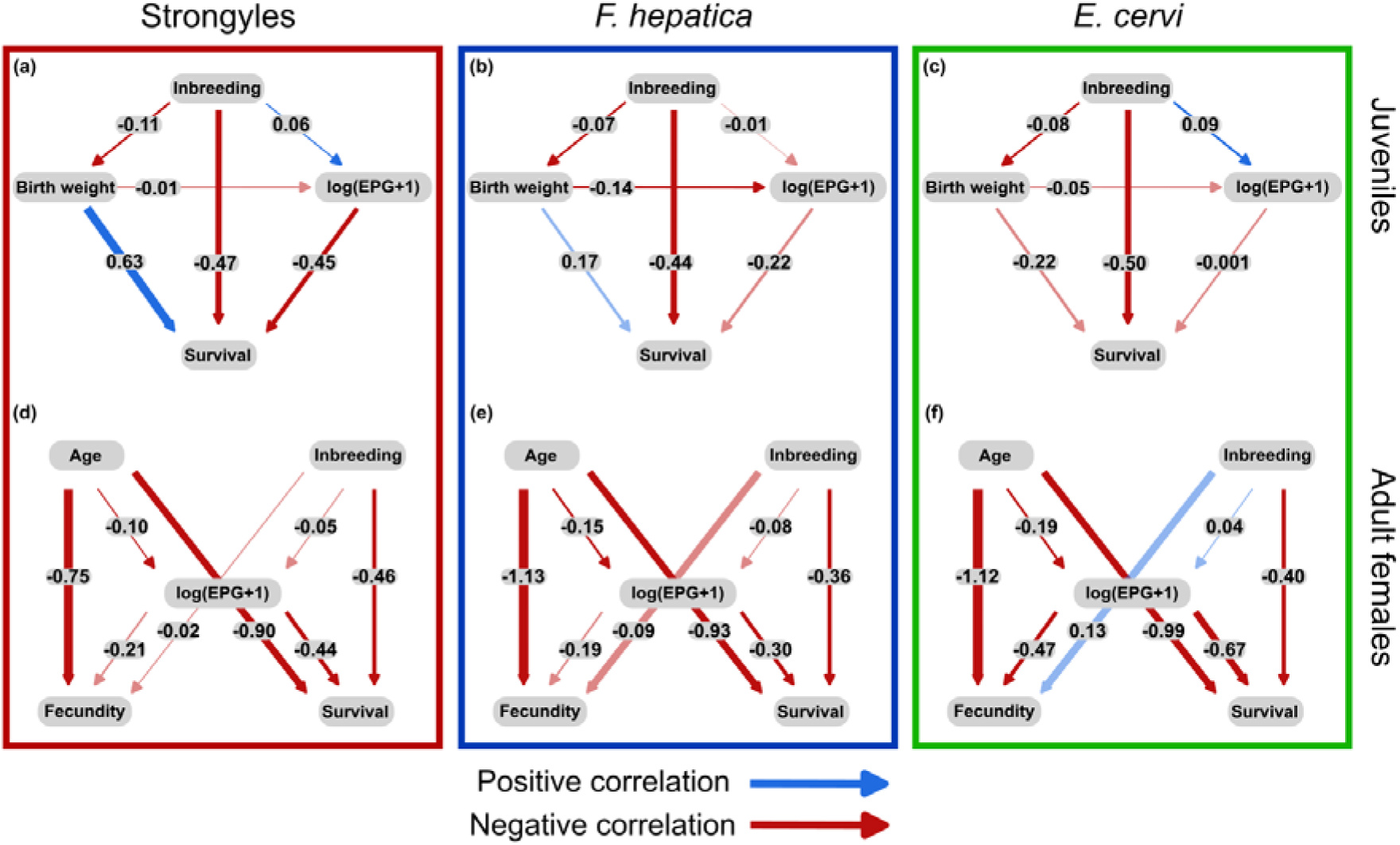
DAGs denoting the relationships between inbreeding, parasitism, and fitness in juvenile red deer for the (a) strongyle, (b) *F. hepatica*, and (c) *E. cervi* datasets as well as the relationships between age, inbreeding, parasitism, and fitness for the (d) strongyle, (e) *F. hepatica*, and (f) *E. cervi* datasets in adult female red deer. Solid lines represent standardized path coefficients of each predictor, which can be interpreted as effect sizes. Blue lines denote positive relationships, while red denote negative relationships. Significance of the effect size is denoted by opaque lines, while non-significant effect sizes are represented by faded lines. Thickness of the lines is proportional to the strength of the effects.

To better visualize these independent paths between inbreeding and juvenile survival in the strongyle dataset, we plotted the direct relationships between F_ROH_ and birth weight, F_ROH_ and strongyle lEPG, F_ROH_ and overwinter survival, strongyle lEPG and overwinter survival, as well as the indirect relationship between F_ROH_ and overwinter survival mediated by strongyles (Fig. 2). Birth weight declined by ∼17% across the range of F_ROH_ (Fig. 2a), while strongyle lEPG increased by ∼20% (Fig. 2b). Overwinter survival was reduced by ∼83% and ∼19% across the ranges of F_ROH_ (Fig. 2c) and strongyle lEPG (Fig. 2d), respectively. Taken together, the indirect, strongyle-mediated effect of F_ROH_ reduced overwinter survival by ∼3% across the range of F_ROH_ (Fig. 2e).

**Figure 2.**
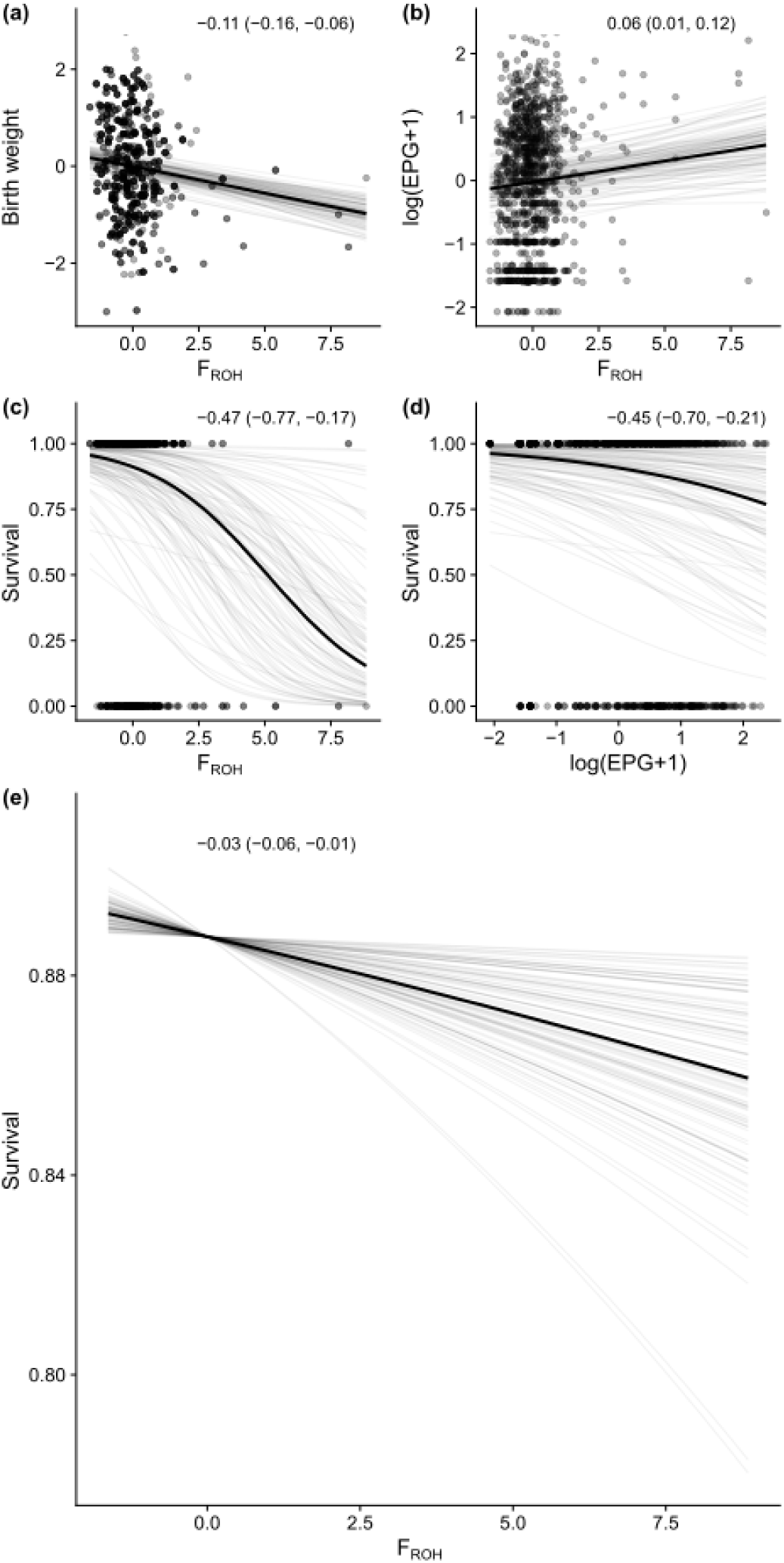
Direct relationships between inbreeding and birth weight (a), inbreeding and strongyle parasitism (b), inbreeding and overwinter survival (c), parasitism and overwinter survival (d), and the indirect, strongyle-mediated relationship between inbreeding and overwinter survival (e). The x-axes denote individual F_ROH_ (a,b,c,e) or individual lEPG (d), with birth weight (a), lEPG (b), or overwinter survival (c-e) on the y-axes. Results are taken from the individual models analyzing the juvenile dataset. The dark black line represents the mean of the posterior distribution for the standardized model estimates, the light grey lines are 100 random draws from the posterior to represent uncertainty. Points denote individual samples, with transparency to allow for visualization of overplotting. The inset text in each panel represents the standardized beta coefficients and associated 95% credible intervals from each regression, which can be interpreted as effect sizes.

### Inbreeding depression across age classes

Our analysis of adult females revealed that the deleterious effects of inbreeding persist beyond the juvenile stage. Specifically, we found that inbreeding moderately reduced overwinter survival in all three datasets, though we did not find any relationships between F_ROH_ and lEPG in any of the datasets (Fig. 1d-f). We also directly linked parasitism and fitness, finding moderate, negative associations between lEPG and survival in the strongyle and *F. hepatica* datasets, in addition to a large negative relationship between lEPG and survival in the *E. cervi* dataset. lEPG was also associated with moderate reductions in fecundity in the *E. cervi* dataset (Fig. 1f). Age was associated with weak declines in lEPG for all three parasites, yet was simultaneously associated with very strong declines in survival and especially fecundity (Fig. 1).

Though it was not significant, we found that more inbred females had fewer strongyles on average (Fig. 1d). Inbreeding increased strongyle parasitism in juveniles, with such increased parasitism resulting in increased juvenile mortality (Fig. 1a). These conflicting patterns across age classes suggest that the combined effects of inbreeding and parasitism may be removing highly inbred as well as highly parasitized individuals via within-generation purging, thus explaining the lack of a pattern in adults. Indeed, the extreme values of both F_ROH_ and strongyle lEPG values are greatly reduced in adult females compared to that of juveniles (Fig. S1).

## Discussion

Identifying the mechanisms by which inbreeding depression manifests is a key first step in moving beyond a phenomenological understanding of the consequences of inbreeding, and crucial to this is understanding if and how parasites mediate inbreeding depression in wild populations. Here, using data from an exceptionally well-characterized ungulate system we linked the deleterious effects of inbreeding to parasitism, and then connected both of these negative factors to host survival in juveniles and adult females. We found that more inbred juvenile deer experienced an increased burden of strongyle nematodes, and increases in strongyle lEPG resulted in decreased overwinter survival – separately from two independent negative correlations between inbreeding and overwinter survival (one direct relationship and one indirect relationship via birth weight). These findings not only support previous findings in this system (22, 25, 28, 29), but also serve as only the second example of parasite-mediated inbreeding depression in a wild population.

The only other study to explicitly link inbreeding to parasite burden, with a subsequent link through to fitness, was Coltman et al. (24). Like our study, Coltman et al. leveraged data from a long-term study of a wild ungulate, Soay sheep (*Ovis aries*). Similar to our results, they found that individual-level inbreeding resulted in increased strongyle burden, with strongyles being negatively associated with overwinter survival in both adults and juveniles (24). A key advance between Coltman et al. and ours is the use of a genome-wide measure of inbreeding, F_ROH_. Coltman et al. use heterozygosity from 14 microsatellite loci, which is an imprecise representation of genome-wide IBD.

Despite finding strong, negative associations between strongyle lEPG and juvenile survival, we did not find a relationship between this measure of fitness and parasitism by *F. hepatica* or *E. cervi* FECs and survival. Previous work has shown that increasing counts of both strongyles and liver fluke were associated with reduced survival in juvenile red deer (25), so it is interesting that we did not find a link with *F. hepatica*. The cumulative nature of our analyses may provide an explanation. Specifically, Acerini et al. (25) found that only summer *F. hepatica* counts in yearlings were negatively associated with reduced overwinter survival. Here, we had more years of data (thus a larger sample size) and included all parasite counts from a year and analyzed calves and juveniles together to gain a more general understanding of the relationships between parasitism and inbreeding and not overfit our models, thus we may have lost some of the nuance of such season- and age-class-specific tests.

The strength of the direct relationship between strongyles and survival is similar to that of the direct relationship between inbreeding and survival, and these relationships are independent of one another. Further, we showed that F_ROH_ was negatively associated with birth weight in the strongyle dataset, mirroring the finding of Hewett et al. (28). However, the remaining and strong negative relationship between F_ROH_ and survival points to additional, untested routes. That is, it could be that inbreeding has negative effects on other fitness-associated variables, and these factors provide further mechanisms by which inbreeding depression operates within this system. For example, inbreeding experiments with *Drosophila simulans* showed that inbreeding reduced development time (30), and early development is a key predictor of juvenile survival in this population of red deer (31, 32). Birth weight, birth day, and parasitism were already included in our analyses, but other aspects of early development may have been negatively impacted by inbreeding and resulted in the observed, parasite-independent increase in overwinter mortality among inbred juveniles. In the red deer on Rum, such effects of early development can extend beyond the juvenile stage to affect adult reproduction and size (33). Our finding that adult female survival decreased with inbreeding, independent of the effects of age and parasitism, confirms previous findings that the effects of inbreeding continue past the juvenile stage (22, 28).

This study has implications for the conservation of endangered populations. It has long been recognized that lack of genetic variation and/or the fixation of deleterious recessive alleles may make populations susceptible to disease. Two examples where this link has been suggested in the past are a cheetah (*Acinonyx jubatus*) colony that was decimated by a coronavirus in Oregon in 1983 (34) and the extreme susceptibility of Père David’s deer (*Elaphurus davidianus*) to malignant catarrhal fever when imported to New Zealand (35). This link is supported by laboratory experiments. For example, extremely inbred (i.e., pedigree inbreeding coefficients of ∼0.25 - 1) *Drosophila melanogaster* were more susceptible to infection by *Serratia marcesans* bacteria and this increased susceptibility was due to reduced resistance to infection in more inbred individuals (36). The red deer of Rum, with mean F_ROH_ = 0.06, are not especially inbred compared with some of the other species of conservation concern which have now been assessed by F_ROH_. Although the depth of genomic information and minimum ROH calls complicate comparison, populations of killer whales (*Orca orca*) (37), tigers (*Panthera tigris*) (38), Eurasian lynx (*Lynx lynx*) (39), Scandinavian wolves (*Canis lupus*) (40), and European ibex (*Capra ibex*) (41) have all been recorded with higher mean F_ROH_ values than the Rum red deer. If we are able to identify an infection route for inbreeding depression in the Rum red deer, it seems likely such effects exist in these other populations and may be even stronger. Future study of parasite-mediated inbreeding depression using methods similar to ours offer a non-invasive way to quantify how vulnerable wild populations are to disease risk.

In conclusion, our study has found clear evidence for parasite-mediated inbreeding depression. Parasitism is a ubiquitous species interaction affecting host fitness, host species interactions, and host evolution across the Tree of Life (5, 42–44). Through the use of high-quality, individual-level data on variation in parasitism and inbreeding, our study has identified parasites as a mechanism by which inbreeding depression manifests in wild populations. While there is more work needed to understand parasite-mediated inbreeding depression, this is a crucial first step. Genomic estimators of inbreeding are more available now than ever, and observational approaches are less invasive, which is important for endangered or threatened populations. Approaches like those utilized in this study could therefore be helpful for better understanding inbreeding depression in the wild. Additionally, more studies of other host-parasite systems are needed to increase our knowledge of if and how a common and widespread species interaction acts to drive inbreeding depression in wild populations of hosts in natural settings, as we know little about the physiological mechanisms inbreeding depression acts through, and the idea that selection by parasites might be involved is very much under-studied. Such studies could be useful and provide crucial information to conservation managers and others working with small and threatened populations that are unable to conduct more invasive sampling or manipulations.

## Methods

### Study system

We collected data from a focal host population of red deer on the north block of the Isle of Rum, Scotland (57°N,6°20’W). The study area runs ∼4km north to south and ∼3km east to west with a total area of ∼12.7km^2^. The population of deer within the study area is wild, unmanaged, and free from both predation and hunting. ∼90% of calves born in the study area are caught soon after birth and weighed, permanently marked, and sampled for genetic analysis; individuals unsampled at this stage are recognized from natural features and are often sampled later via darting, cast antlers, or *post mortem*.

While the deer have been studied in some capacity since 1972 (45), data on the helminth parasite burden of the population has been non-invasively collected since 2016 by collecting fecal samples three times a year in April (Spring), August (Summer), and November (Autumn). Briefly, observers note individually-recognized deer defecating from a distance and collect the fecal samples without disturbing the deer. Samples are then placed into plastic bags to keep the samples as anaerobic as possible and refrigerated at 4°C to prevent hatching or development of parasite propagules, with subsequent parasitological examination being conducted within three weeks in the case of strongyles(46). Detailed methods can be found in Albery et al. (46).

For our analysis of parasitism we focused on three of the most common parasite taxa infecting the red deer: strongyles, *F. hepatica*, and *E. cervi*. Strongyles have a direct lifecycle in which infective stages contaminate vegetation via fecal pellets and are subsequently consumed by a new host (47). *F. hepatica* (47) and *E. cervi* (48) both have indirect lifecycles involving a snail intermediate host (the dwarf pond snail *Galba truncatula* and a number of terrestrial snails and slugs, respectively). After infecting and emerging from their intermediate hosts, larval *F. hepatica* contaminate vegetation near water bodies which are consumed by the deer final host. In contrast, deer become infected with *E. cervi* by consuming the intermediate snail host itself. While strongyle infections develop quickly such that calves excrete eggs within 2-3 months of birth, *F. hepatica* and *E. cervi* have longer prepatent periods, resulting in low prevalences of *F. hepatica* and *E. cervi* for juveniles relative to adults (46, 49, 50) (Fig. S2).

Individual inbreeding was estimated using the runs of homozygosity (ROH)-based inbreeding coefficient, F_ROH,_ which captures realized IBD (22). All sampled deer have been genotyped on the Illumina cervine 50K Single Nucleotide Polymorphism (SNP) and after QC 37,396 SNPs were used to search for ROHs of minimum length 2.5Mb (hereafter F_ROH2.5MB_) in PLINK v2.0 using the physical locations of SNPs from the red deer genome assembly mCerEla1.1 (51). For details on QC and ROH searching see (52). With these conditions, mean F_ROH2.5MB_ at birth in the Rum deer is 0.06 and ranges from 0 – 0.35 (52) (Fig. S3).

In juveniles, we focused on two fitness-associated measures: birth weight and overwinter survival. Birth weight is adjusted from capture weight to account for postnatal growth. Overwinter survival is a binary response, with 1 for calves/yearlings that lived through the winter into the start of the next “deer year”, a period running from May 1 – June 30, and 0 for those calves/yearlings that died before the start of the next deer year. These data included all calves and yearlings that we had parasite, immunity, and inbreeding data for. Individuals that were shot as juveniles when they roamed outside of the study area were removed from the analysis, as we were only interested in natural mortality.

For female fitness we focused on overwinter survival and fecundity. Overwinter survival was calculated as above, while fecundity was calculated as a binary response, with 1 for females that produced a calf in the next deer year, and 0 for those females that did not produce a calf in the next deer year. As with juveniles, females that were shot when they ventured outside of the study area were removed from the analysis.

### Statistical analyses

Parasite fecal egg counts (FECs) and F_ROH2.5MB_ values varied across space in both the juvenile and adult datasets (Fig. S4). Confirming results from previous studies of this system (50, 53) we found that strongyle and *E. cervi* FECs in both datasets were more abundant in the north of the study area, while *F. hepatica* was more abundant towards the middle of the study area in the adult females (Fig. S4). Relatively high values of F_ROH_ in the juveniles were found in multiple locations throughout the study area, while they tended to be concentrated in the north for the adult females (Fig. S4). To control for this spatial autocorrelation, we used Integrated Nested Laplace Approximation (INLA) models for all analyses. INLA models are a deterministic Bayesian approach which allow for the quantification of spatial effects and have been increasingly used for spatial analyses (53–55). To calculate annual centroids for each individual we utilized the census data, where individuals’ identities and locations (to the nearest 100m) were recorded. We then calculated the annual centroid using a previously described pipeline for this population (56, 57) using all observations of each individual in each year. This approach uses a kernel density estimator, taking individuals’ annual centroids and fitting a two-dimensional smooth to the distribution of the data, producing a two-dimensional spatial distribution of the population. We fit all models in R version 4.2.2 (58) using the R-INLA package (59, 60).

### Model construction

We constructed separate models for each of the three parasites using parasite-specific datasets. For the juveniles the strongyle dataset included all calves and yearlings for which we had strongyle data (*n* = 1,068 records from *n* = 348 deer). The *F. hepatica* dataset included all calves and yearlings for which we had liver fluke data (*n* = 655 records from *n* = 301 deer), excluding data collected from calves in their first summer as *F. hepatica* are prepatent this early in a calf’s life (46, 49). Finally, the *E. cervi* dataset included all calves and yearlings for which we had tissue worm data (*n* = 419 records from *n* = 213 deer), excluding data collected from calves in their first summer and autumn as *E. cervi* have an even longer prepatent period (46, 49). For adult females the strongyle dataset included *n* = 1,698 records from *n* = 170 deer, the *F. hepatica* dataset included *n* = 1,393 records from *n* = 168 deer, and the *E. cervi* dataset included *n* = 1,313 records from *n* = 163 deer.

To investigate links between inbreeding and parasitism, inbreeding and fitness, and parasitism and fitness, we utilized path analyses with the D-separation method (27), where some variables appear as both fixed effects and response variables (61). While path analyses are a useful tool and allow for tests of hypothesized, “causal” relationships using large amounts of observational data (27, 62, 63), such models estimate correlational relationships and cannot show causality (61). Combining multiple models in this way allows for identification of variables/mediating factors that are driving the overall relationship (27). Specifically, we constructed directed acyclic graphs (DAGs) for each parasite and age class to test if and how a given parasite mediated inbreeding depression in this system.

For juveniles, each DAG was composed of three individual models quantifying the relationships between inbreeding, parasitism, and fitness. See Fig. S5 for a graphical representation of these hypothesized relationships. For the first model we used lEPG as our response variable with F_ROH2.5MB_, age category (calf or yearling, accounting for age-category-specific differences (46, 50)), sex (accounting for sex-specific differences (50)), birth weight (in kilograms), and birth day (Julian day, accounting for the fact that individuals born earlier have higher survival (64)) as fixed effects in a Gaussian mixed-effect model with year (categorical) and maternal ID as random effects to account for the correlational structure of the data and maternal effects, respectively. Importantly for this model we mean-centered lEPG values by season to limit collinearity with other fixed effects, thus controlling for seasonal variation in parasite load (46, 50) without including season as a term in our models. For the second model we used birth weight as the response variable with F_ROH2.5MB_, sex, and birth day as fixed effects in a Gaussian mixed-effect model with year and maternal ID as random effects. For the third model we used overwinter survival as the response variable with F_ROH2.5MB_, birth weight, age category, sex, birth day, and lEPG as fixed effects in a mixed-effect logistic regression with year and maternal ID as random effects.

For the adult females, each DAG was similarly composed of three individual models using different model structures (see Fig. S5). For the first model we used lEPG as our response variable with F_ROH2.5MB_, age (a continuous variable, accounting for age-specific differences in parasite load (26)), and reproductive status in the current year (none: did not have a calf, summer yeld: had a calf but it died over the summer, or winter: had a calf and raised the calf up to or through winter, accounting for known relationships between reproductive investment and parasitism (27, 50)) as fixed effects in a Gaussian mixed-effect model with year (categorical) and individual ID as random effects to account for the correlational structure of the data and repeated measures, respectively. We again mean-centered lEPG values by season. For the second model we used overwinter survival as the response variable with F_ROH2.5MB_, age, lEPG, and reproductive status in the current year as fixed effects in a mixed-effect logistic regression model with year and individual ID as random effects. For the third model we used fecundity (binary: 1 for females that had a calf in the subsequent year and 0 for females that did not) as the response variable in a mixed-effect logistic regression with F_ROH2.5MB_, age, lEPG, and reproductive status in the current year as fixed effects with year and individual ID as random effects.

Connecting these models together using DAGs allows us to investigate the hypothesized causal relationships between inbreeding, parasitism, and fitness, identifying the routes by which inbreeding depression manifests within this population. We scaled all continuous variables to have a mean of 0 and SD of 1. Though each of the models described above included important covariates such as age category, sex, and reproductive status due to their known relationships with parasitism in this system, we only present results for the relationships between inbreeding, parasitism, and fitness. The full model results can be found in the Supplemental Materials (Fig. S6-7).

## Author contributions

AZH & JMP conceived the study. AZH, JMP, and GFA designed the study and analyses. SM, AM, and JMP collected field data and samples, KM and AZH quantified parasite samples, AMH and JMP processed DNA samples, conducted QC, and derived F_ROH2.5MB_ values. AZH performed modelling work and analyzed data. AZH wrote the first draft of the manuscript, and all authors contributed substantially to revisions.

## Acknowledgements

We thank NatureScot for permission to work on the Isle of Rum, those involved in running the project including Tim Clutton-Brock and Loeske Kruuk and the many volunteers and researchers who have helped at the field site during this study. AZH benefitted from the musical inspiration of The Darkness. Genotyping was conducted at the Wellcome Trust Clinical Research Facility Genetics Core. We thank Susan Johnston and Jisca Huisman for additional QC of SNP data. We also thank Fiona Kenyon, David McBean, and others from the Moredun Institute for their help in quantifying FECs.

## References

1. D. Charlesworth, J. H. Willis, The genetics of inbreeding depression. Nature Reviews Genetics 10, 783–796 (2009).

2. C. Darwin, The effects of cross and self fertilization in the vegetable kingdom (John Murray, London, 1876).

3. M. Kardos, G. Luikart, F. W. Allendorf, Measuring individual inbreeding in the age of genomics: marker-based measures are better than pedigrees. Heredity 115, 63–72 (2015).

4. M. Kardos, H. R. Taylor, H. Ellegren, G. Luikart, F. W. Allendorf, Genomics advances the study of inbreeding depression in the wild. Evolutionary Applications 9, 1205–1218 (2016).

5. A. Z. Hasik, A. M. Siepielski, Parasitism shapes selection by drastically reducing host fitness and increasing host fitness variation. Biology Letters 18, 20220323 (2022).

6. W. K. Potts, C. J. Manning, E. K. Wakeland, The role of infectious disease, inbreeding and mating preferences in maintaining MHC genetic diversity: an experimental test. Philosphical Transactions of the Royal Society B: Biological Sciences 346, 19940154 (1994).

7. S. Sommer, The importance of immune gene variability (MHC) in evolutionary ecology and conservation. Frontiers in Zoology 2, 1–18 (2005).

8. S. K. J. R. Auld, S. K. Tinkler, M. C. Tinsley, Sex as a strategy against rapidly evolving parasites. Proceedings of the Royal Society B 283, 20162226 (2016).

9. W. D. Hamilton, R. Axelrod, R. Tanese, Sexual reproduction as an adaptation to resist parasites (a review). Proceedings of the National Academy of Sciences of the United States of America 87, 3566–3573 (1990).

10. D. Ebert, P. D. Fields, Host–parasite co-evolution and its genomic signature. Nature Reviews Genetics 21, 754–768 (2020).

11. C. Eizaguirre, T. L. Lenz, M. Kalbe, M. Milinski, Divergent selection on locally adapted major histocompatibility complex immune genes experimentally proven in the field. Ecology Letters 15, 723–731 (2012).

12. J. Klein, C. O’Huigin, J. Deutsch, MHC polymorphism and parasites. Philosphical Transactions of the Royal Society B: Biological Sciences 346, 19940152 (1994).

13. N. K. Whiteman, K. D. Matson, J. L. Bollmer, P. G. Parker, Disease ecology in the Galápagos Hawk (*Buteo galapagoensis*): host genetic diversity, parasite load and natural antibodies. Proceedings of the Royal Society B 273, 20053396 (2006).

14. E. A. MacDougall-Shackleton, E. P. Derryberry, J. Foufopoulos, A. P. Dobson, T. P. Hahn, Parasite-mediated heterozygote advantage in an outbred songbird population. Biology Letters 1, 20040264 (2005).

15. K. Acevedo-Whitehouse et al., Contrasting effects of heterozygosity on survival and hookworm resistance in California sea lion pups. Molecular Ecology 15, 1973–1982 (2006).

16. J. Slate et al., Understanding the relationship between the inbreeding coefficient and multilocus heterozygosity: theoretical expectations and empirical data. Heredity 93, 255–265 (2004).

17. F. Balloux, W. Amos, T. Coulson, Does heterozygosity estimate inbreeding in real populations? Mol. Ecol. 13, 3021–3031 (2004).

18. R. J. Sardell, L. F. Keller, P. Arcese, T. Bucher, J. M. Reid, Comprehensive paternity assignment: genotype, spatial location and social status in song sparrows, Melospiza Melodia. Mol. Ecol. 19, 4352–4364 (2010).

19. J. I. Hoffman et al., High-throughput sequencing reveals inbreeding depression in a natural population. Proceedings of the National Academy of Sciences of the United States of America 111, 3775–3780 (2014).

20. J. Yang, S. H. Lee, M. E. Goddard, P. M. Visscher, GCTA: A tool for genome-wide complex trait analysis. Am. J. Hum. Genet. 88, 76–82 (2011).

21. R. McQuillan et al., Runs of homozygosity in European populations. Am. J. Hum. Genet. 83, 359–372 (2008).

22. J. Huisman, L. E. B. Kruuk, P. A. Ellis, T. Clutton-Brock, J. M. Pemberton, Inbreeding depression across the lifespan in a wild mammal population. Proceedings of the National Academy of Sciences 113, 3585–3590 (2016).

23. C. Bérénos, P. A. Ellis, J. G. Pilkington, J. M. Pemberton, Genomic analysis reveals depression due to both individual and maternal inbreeding in a free-living mammal population. Mol. Ecol. 25, 3152–3168 (2016).

24. D. W. Coltman, J. G. Pilkington, J. A. Smith, J. M. Pemberton, Parasite-mediated selection against inbred soay sheep in a free-living island population. Evolution 53, 1259–1267 (1999).

25. C. I. Acerini et al., Helminth parasites are associated with reduced survival probability in young red deer. Parasitology 149, 1702–1708 (2022).

26. G. F. Albery et al., Divergent age-related changes in parasite infection occur independently of behaviour and demography in a wild ungulate. Philosophical Transactions of the Royal Society B 10.1098/rstb.2023.0508 (2024).

27. G. F. Albery et al., Fitness costs of parasites explain multiple life-history trade-offs in a wild mammal. The American Naturalist 197, 324–335 (2021).

28. A. M. Hewett, S. E. Johnston, A. Morris, S. Morris, J. M. Pemberton, Genetic architecture of inbreeding depression may explain its persistence in a population of wild red deer. Molecular Ecology 33, e17335 (2024).

29. C. A. Walling et al., Inbreeding depression in red deer calves. BMC Evolutionary Biology 11, 1–13 (2011).

30. L. I. Wright, T. Tregenza, D. J. Hosken, Inbreeding, inbreeding depression and extinction. Conservation Genetics 9, 833–843 (2007).

31. T. H. Clutton-Brock, M. Major, S. D. Albon, F. E. Guiness, Early development and population dynamics in red deer. I. Density-dependent effects on juvenile survival. Journal of Animal Ecology 56, 53–67 (1987).

32. L. E. B. Kruuk, T. H. Clutton-Brock, K. E. Rose, F. E. Guinness, Early determinants of lifetime reproductive success differ between the sexes in red deer. Proceedings of the Royal Society B 266, 19990828 (1999).

33. S. D. Albon, T. H. Clutton-Brock, F. E. Guiness, Early development and population dynamics in red deer. II. Density-independent effects and cohort variation. Journal of Animal Ecology 56, 69–81 (1987).

34. S. J. O’Brien et al., Genetic basis for species vulnerability in the cheetah. Science 227, 1428–1434 (1985).

35. M. B. Orr, C. G. Mackintosh, An outbreak of malignant catarrhal fever in Père David’s deer (*Elaphurus davidianus*). New Zealand Veterinary Journal 36, 19–21 (1988).

36. D. Spielman, B. W. Brook, D. A. Briscoe, R. Frankham, Does inbreeding and loss of genetic diversity decrease disease resistance? Conservation Genetics 5, 439–448 (2004).

37. M. Kardos et al., Inbreeding depression explains killer whale population dynamics. Nature Ecology & Evolution 7, 675–686 (2023).

38. A. Khan et al., Genomic evidence for inbreeding depression and purging of deleterious genetic variation in Indian tigers. Proceedings of the National Academy of Sciences 118, e2023018118 (2021).

39. F. Abascal et al., Extreme genomic erosion after recurrent demographic bottlenecks in the highly endangered Iberian lynx. Genome Biology 17, 1–19 (2016).

40. M. Kardos et al., Genomic consequences of intensive inbreeding in an isolated wolf population. Nature Ecology & Evolution 2, 124–131 (2017).

41. C. Grossen, F. Guillaume, L. F. Keller, D. Croll, Purging of highly deleterious mutations through severe bottlenecks in Alpine ibex. Nature Communications 11, 1–12 (2020).

42. A. Buckling, D. J. Hodgson, ShortlJterm rates of parasite evolution predict the evolution of host diversity. Journal of Evolutionary Biology 20, 1682–1688 (2007).

43. A. Betts, C. Gray, M. Zelek, R. C. MacLean, K. C. King, High parasite diversity accelerates host adaptation and diversification. Science 360, 907–911 (2018).

44. A. Z. Hasik et al., Resetting our expectations for parasites and their effects on species interactions: a meta-analysis. Ecology Letters 10.1111/ele.14139, 184–199 (2023).

45. J. M. Pemberton, L. E. B. Kruuk, T. Clutton-Brock, The unusual value of long-term studies of individuals: the example of the Isle of Rum Red Deer Project. Annual Review of Ecology, Evolution, and Systematics 53, 327–351 (2022).

46. G. F. Albery et al., Seasonality of helminth infection in wild red deer varies between individuals and between parasite taxa. Parasitology 145, 1410–1420 (2018).

47. M. A. Taylor, R. L. Coop, R. L. Wall, “Parasites of ungulates” in Veterinary parasitology M. A. Taylor, R. L. Coop, R. L. Wall, Eds. (John Wiley & Sons, Ltd, Hoboken, New Jersey, 2016), pp. 761–815.

48. P. C. Mason, *Elaphostrongylus cervi* - a review. Surveillance 16, 3–10 (1989).

49. A. A. Gajadhar, S. V. Tessaro, W. D. Yates, Diagnosis of *Elaphostrongylus cervi* infection in New Zealand red deer (*Cervus elaphus*) quarantined in Canada, and experimental determination of a new extended prepatent period. The Canadian Veterinary Journal 35, 433–437 (1994).

50. A. Z. Hasik et al., Population density drives increased parasitism via greater exposure and reduced resource availability in wild red deer. Authorea 10.22541/au.173143117.74697497/v1 (2024).

51. J. M. Pemberton, S. E. Johnston, T. J. Fletcher, D. T. o. L. Consortium, The genome sequence of the red deer, *Cervus elaphus* Linnaeus 1758. Wellcome Open Research 6, 336 (2021).

52. A. M. Hewett, M. A. Stoffel, L. Peters, S. E. Johnston, J. M. Pemberton, Selection, recombination and population history effects on runs of homozygosity (ROH) in wild red deer (*Cervus elaphus*). Heredity 130, 242–250 (2023).

53. G. F. Albery, D. J. Becker, F. Kenyon, D. H. Nussey, J. M. Pemberton, The fine-scale landscape of immunity and parasitism in a wild ungulate population. Integrative and Comparative Biology 59, 1165–1175 (2019).

54. G. F. Albery et al., Negative density-dependent parasitism in a group-living carnivore. Proceedings of the Royal Society B 287, 20202655 (2020).

55. A. F. Zuur, E. N. Ieno, A. A. Saveliev, Beginner’s guide to spatial, temporal, and sptial-temporal ecological data analydid with R-INLA (Highstat Ltd., Newburgh, Scotland, 2017).

56. G. F. Albery et al., Ageing red deer alter their spatial behaviour and become less social. Nature Ecology & Evolution 6, 1231–1238 (2022).

57. G. F. Albery et al., Multiple spatial behaviours govern social network positions in a wild ungulate. Ecology Letters 24, 676–686 (2021).

58. R. C. Team (2021) R: A language and environment for statistical computing. (RStudio, Inc., Vienna, Austria).

59. T. G. Martins, D. Simpson, F. Lindgren, H. Rue, Bayesian computing with INLA: new features. Computational Statistics & Data Analysis 67, 68–83 (2013).

60. H. Rue, S. Martino, N. Chopin, Approximate Bayesian Inference for Latent Gaussian models by using Integrated Nested Laplace Approximations. Journal of the Royal Statistical Society Series B: Statistical Methodology 71, 319–392 (2009).

61. B. Shipley, Confirmatory path analysis in a generalized multilevel context. Ecology 90, 363–368 (2009).

62. A. Hasik, J. Bried, D. Bolnick, A. Siepielski, Is the local environment more important than within-host interactions in determining coinfection?

63. A. Z. Hasik, A. M. Siepielski, A role for the local environment in driving specieslJspecific parasitism in a multilJhost parasite system. Freshwater Biology 67, 1571–1583 (2022).

64. T. H. Clutton-Brock, F. E. Guinness, S. D. Albon, Red deer: behavior and ecology of two sexes (University of Chicago Press, Chicago, IL, 1982).

