## Supplementary Information for "Parasite-mediated inbreeding depression in wild red deer"

**Figures S1-S7**

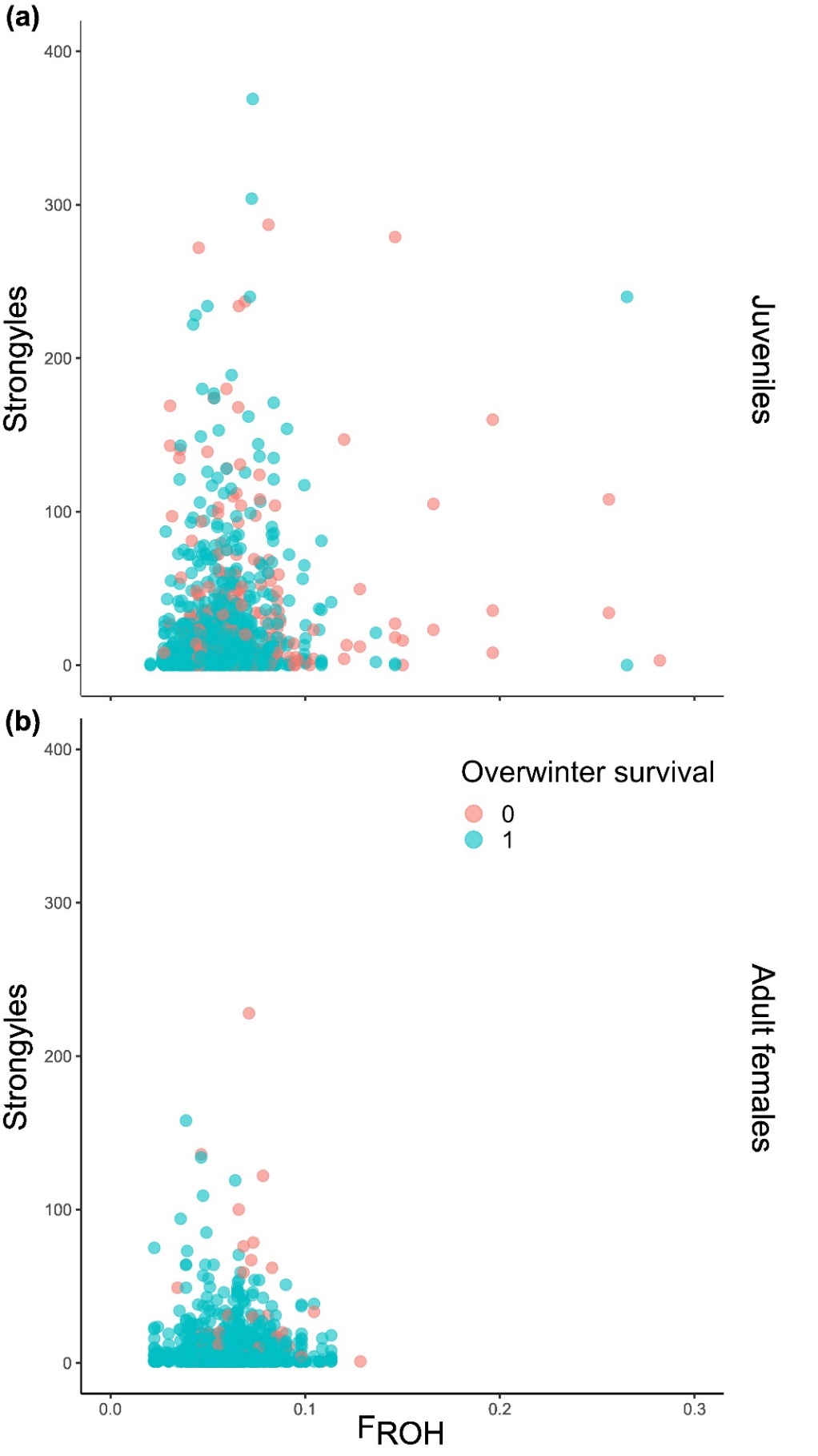

**Figure S1** – Relationships between inbreeding and strongyle parasitism in juveniles (top) and adult females (bottom). The x-axes denote individual F_ROH_ with strongyle fecal egg counts on the y-axes. Points denote raw data values, not accounting for repeated measures, season, age class (juveniles), sex (juveniles), age (adult females), or reproductive status (adult females). Color of points denote overwinter survival.

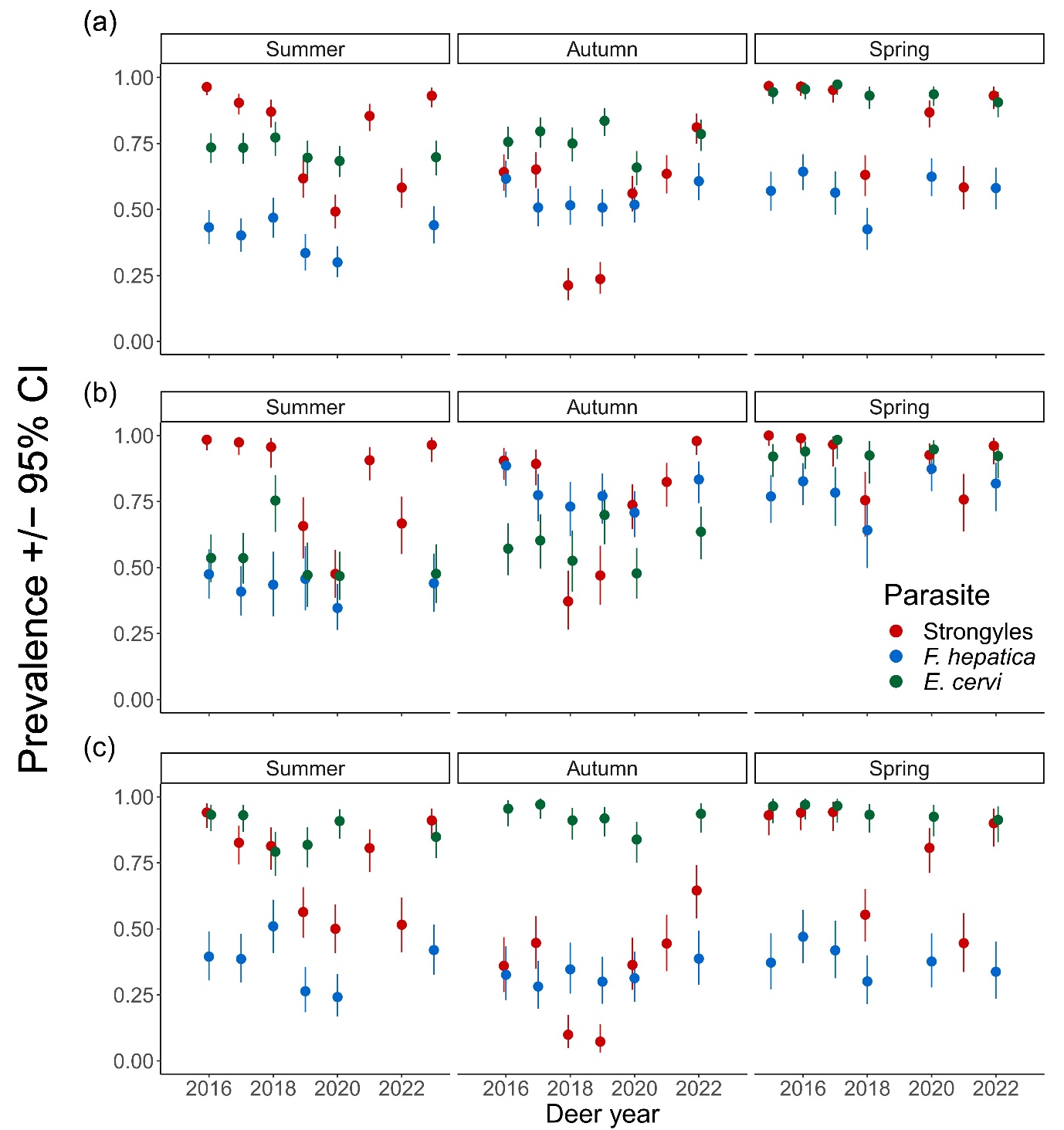
**Figure S2 -** Plots of the seasonal prevalence patterns for each parasite for (a) all deer, (b) juveniles only, and (c) adult females only, with Deer year on the x-axis. The deer year runs from May 1^st^ to April 30^th^, so deer year 2016 ran from May 1^st^ 2016, with first sampling session in summer 2016, to April 30^th^ 2017. Points represent mean prevalence values, error bars denote 95% confidence intervals, and color denotes the parasite.

**Figure S3 –**
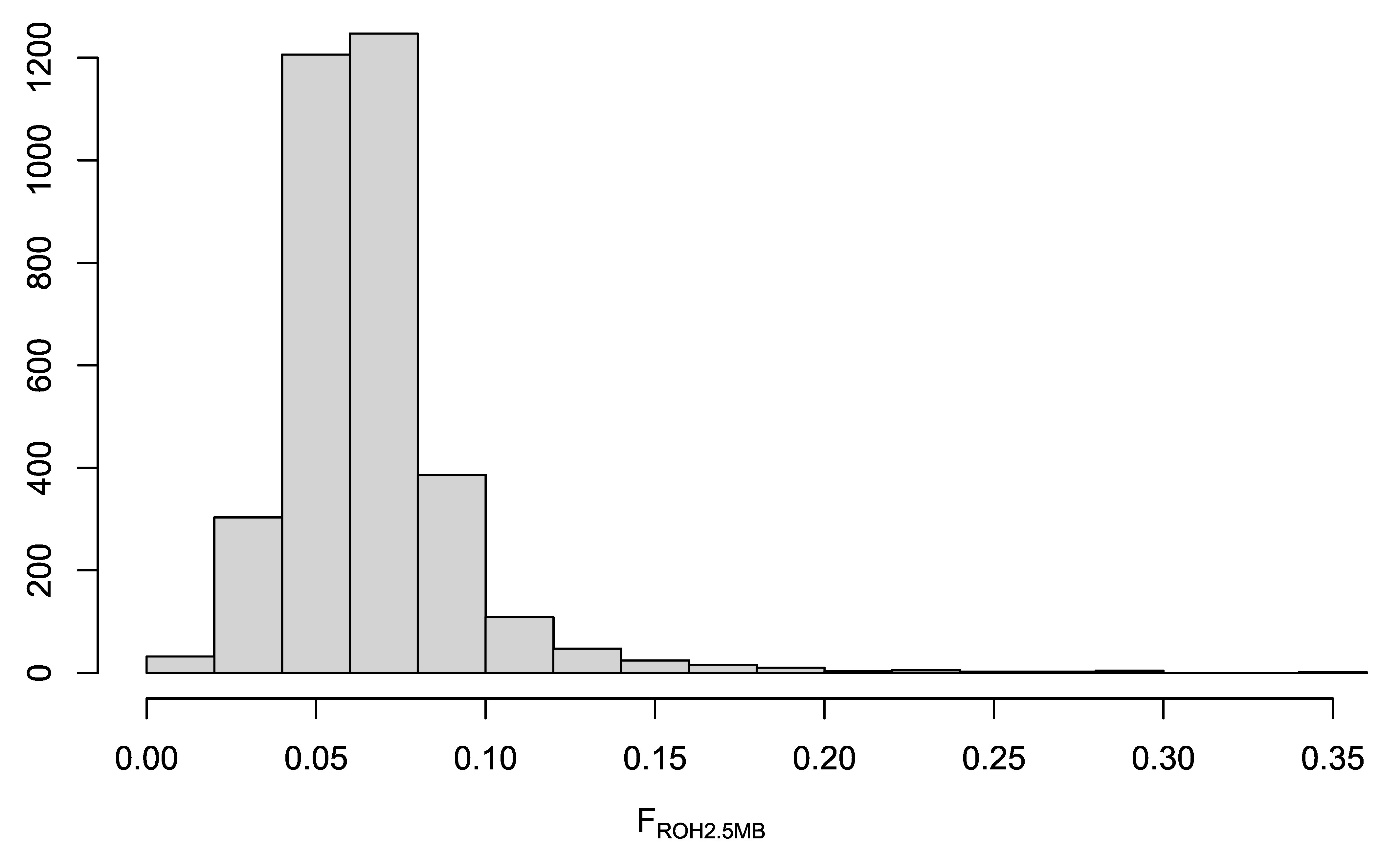
Histogram of the distribution of F_ROH2.5MB_ values for all deer genotyped from the Isle of Rum study population.

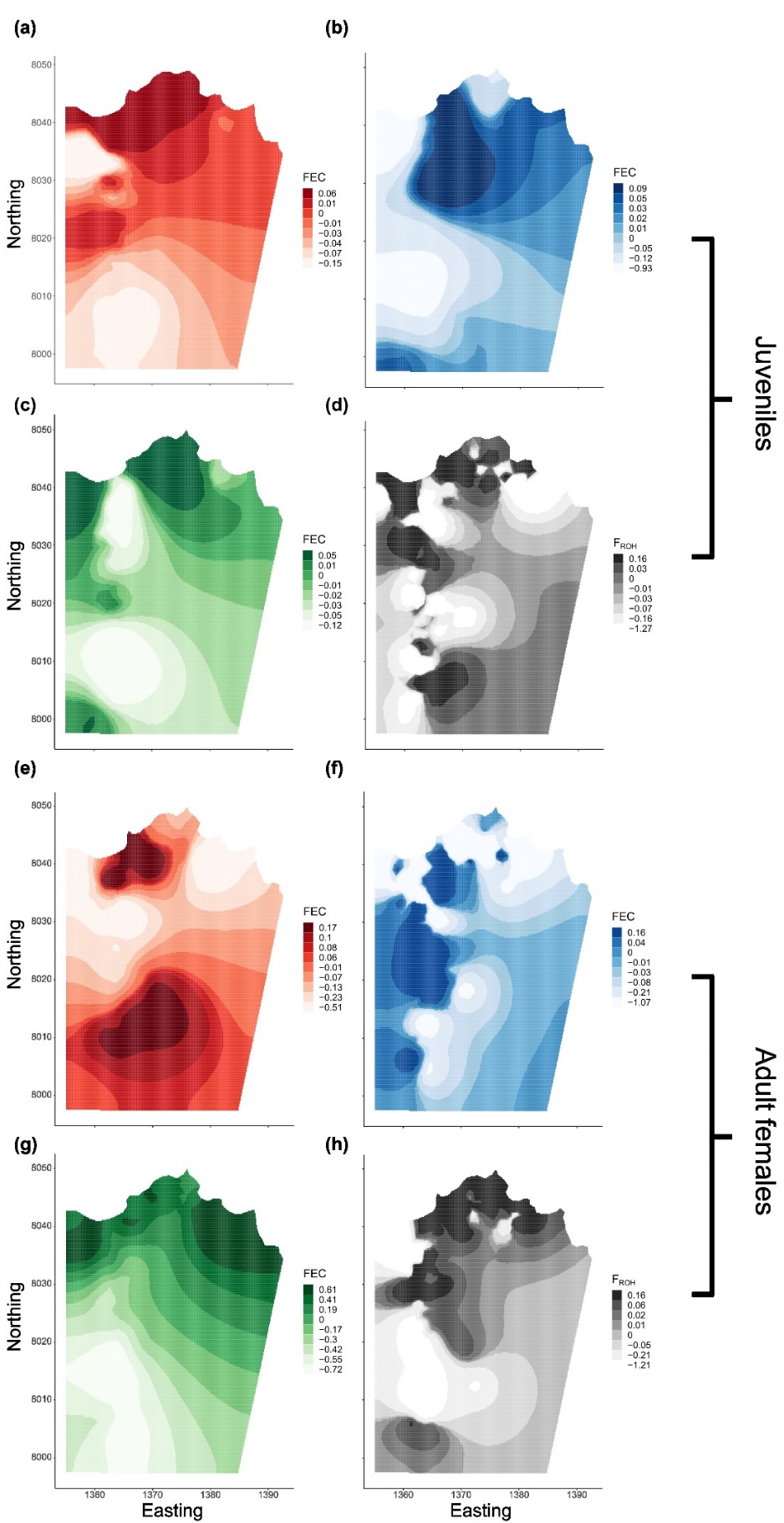

**Figure S4** – Spatial distribution of parasite counts (FEC) for strongyle (a,e), *F. hepatica* (b,f), and *E. cervi* (c,g) and F_ROH_ values (d, h) throughout the study area. (a-d) shows the spatial distributions for juvenile deer, while (e-h) shows the same for adult females. Shown in all plots are projections of the spatially-distributed random effect from the null model for each parasite (a-c and e-g, providing a representation of where parasite counts are higher) and for F_ROH_ (d and h, providing a representation of where inbreeding is the most severe). Shading of the map denotes the lower bounds of quantiles of the spatial effects on the link scale, rounded to two decimal places, with darker colors representing higher parasite FECs (a-c and e-g) or F_ROH_ values (d, h). Easting and Northing are in units of 100m grid squares, with 10 units equaling 1 km. The river at the base of the valley runs along the 1363 Easting.

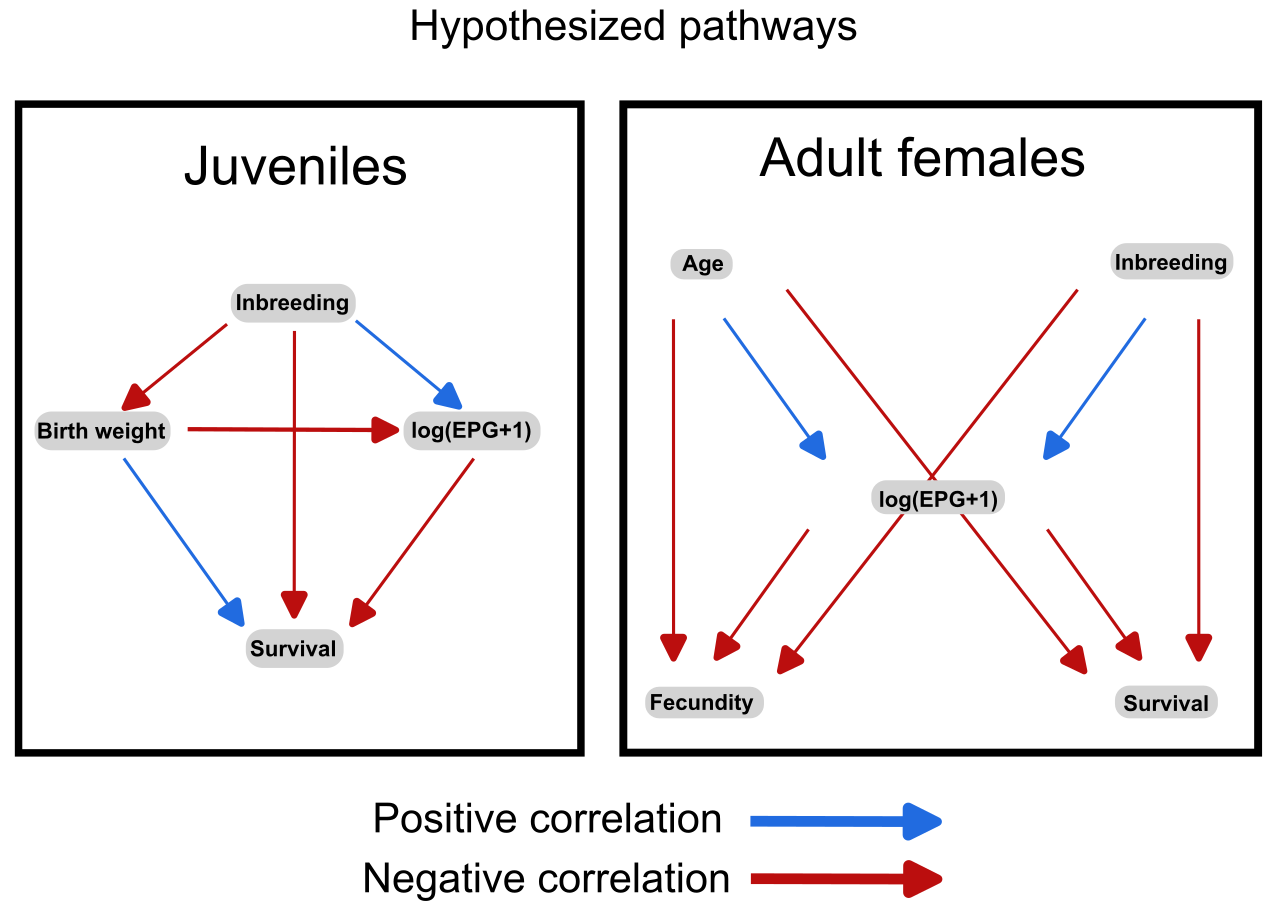
**Figure S5 –** Representation of the hypothesized relationships between inbreeding, parasitism, and fitness in juvenile red deer, in addition to hypothesized relationships between age, inbreeding, parasitism, and fitness in adult female red deer. Effects in the juvenile box are predicted to flow from the top of the plot to the bottom, with inbreeding affecting birth weight, parasitism, and survival, and parasitism and birth weight affecting survival. Effects in the adult female box are also predicted to flow from the top of the plot to the bottom, with both age and inbreeding affecting parasitism and parasitism, age, and inbreeding affecting fecundity and survival. Note that although we predict a positive relationship between age and parasitism in the adult females, this is only predicted for the strongyle dataset, negative relationships are expected in the *F. hepatica* and *E. cervi* datasets due to the observed patterns in prior studies of this system. Positive predicted relationships are shown in blue, while negative relationships are shown in red.

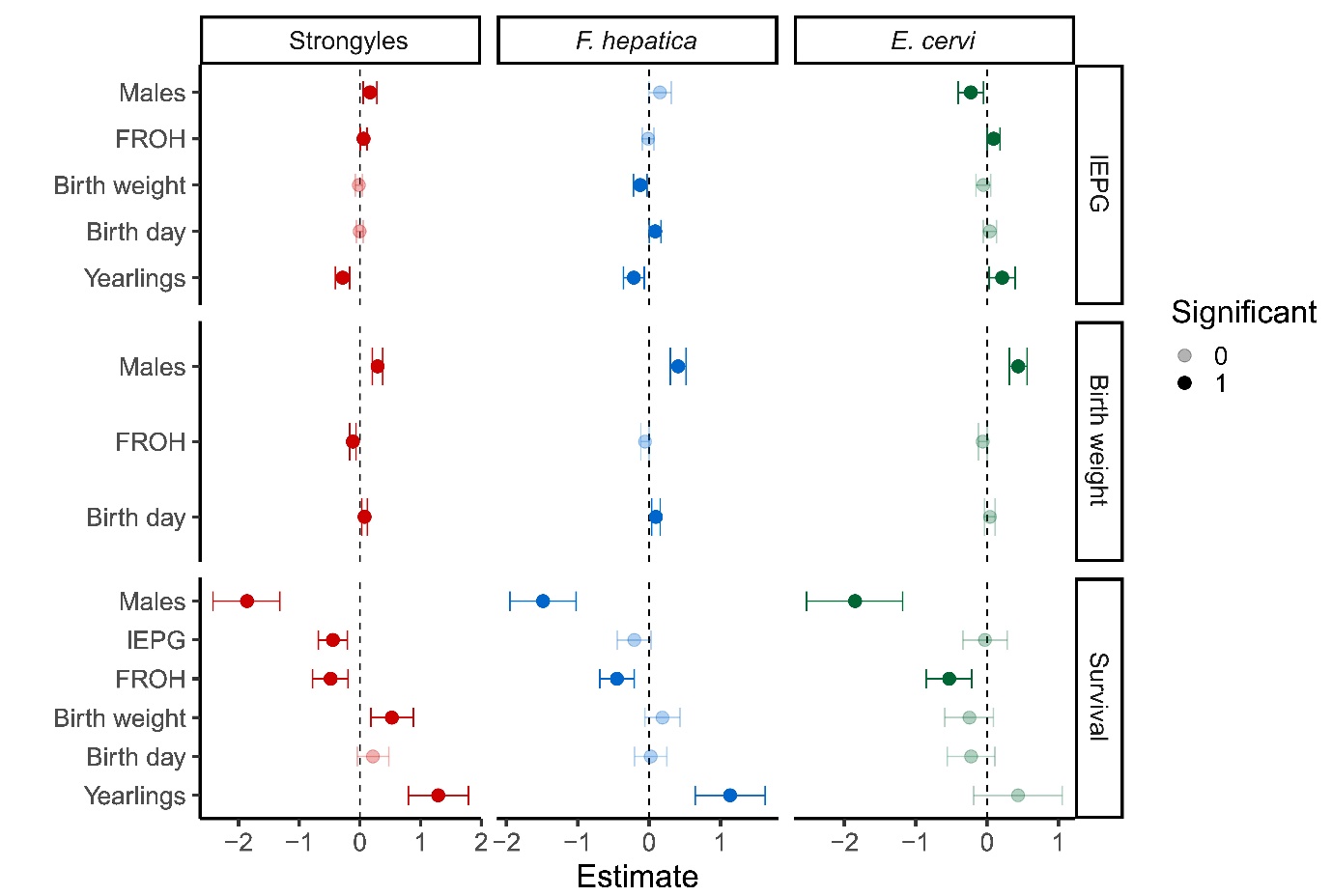
**Figure S6 -** Forest plot representing the full model results from each component of our juvenile SEMs, with the parasite model in the top row, the birth weight model in the middle row, the survival model in the bottom row, and with panels for each parasite. Points represent posterior estimates for mean effect sizes, error bars denote 95% credible intervals in standard deviations, and color denotes the parasite taxa. Significance of the effect size is denoted by the shading of the points

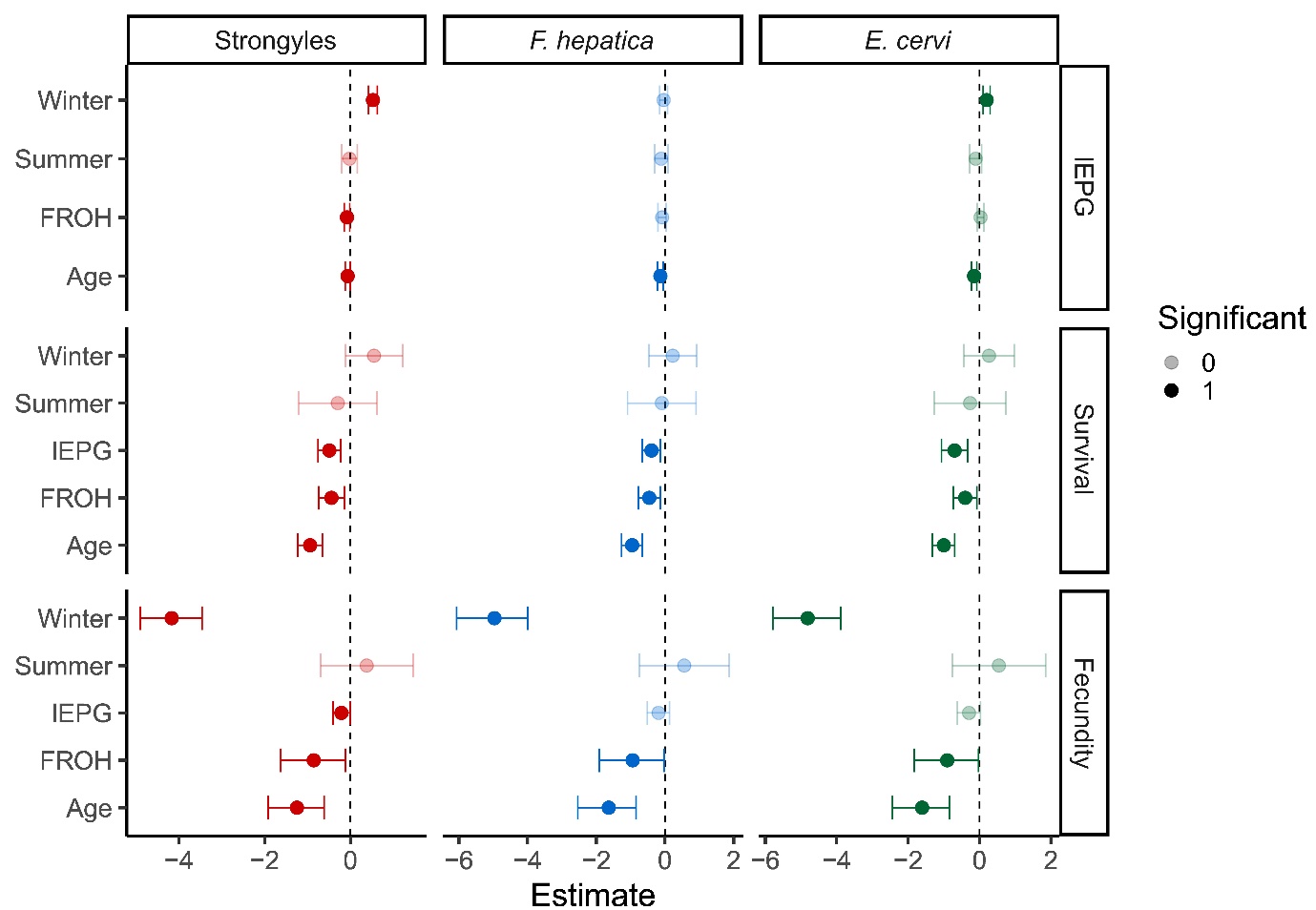
**Figure S7 -** Forest plot representing the full model results from each component of our adult female SEMs, with the parasite model in the top row, the survival model in the middle row, the fecundity model in the bottom row, and with panels for each parasite. Points represent posterior estimates for mean effect sizes, error bars denote 95% credible intervals in standard deviations, and color denotes the parasite taxa. Significance of the effect size is denoted by the shading of the points.
